# Curatr: a web application for creating, curating, and sharing a mass spectral library

**DOI:** 10.1101/170571

**Authors:** Andrew Palmer, Prasad Phapale, Dominik Fay, Theodore Alexandrov

## Abstract

**Motivation:** Identification from metabolomics mass spectrometry experiments requires comparison of fragmentation spectra from experimental samples to spectra from analytical standards. As the quality of identification depends directly on the quality of the reference spectra, manual curation is routine during the selection of reference spectra to include in a spectral library. Whilst building our own in-house spectral library we realised that there is currently no vendor neutral open access tool for for facilitating manual curation of spectra from raw LC-MS data into a custom spectral library.

**Results:** We developed a web application *curatr* for the rapid generation of high quality mass spectral fragmentation libraries for liquid-chromatography mass spectrometry analysis. Curatr handles datasets from single or multiplexed standards, automatically extracting chromatographic profiles and potential fragmentation spectra for multiple adducts. These are presented through an intuitive interface for manual curation before being documented in a custom spectral library. Searchable molecular information and the providence of each standard is stored along with metadata on the experimental protocol. Curatr support the export of spectral libraries in several standard formats for easy use with third party software or submission to community databases, maximising the return on investment for these costly measurements. We demonstrate the use of curatr to generate the EMBL Metabolomics Core Facility spectral library which is publicly available at http://curatr.mcf.embl.de.

**Availability:** The source code is freely available at http://github.com/alexandrovteam/curatr/ along with example data.

**Supplementary information:** A step-by step user manual is available in the supplementary information

## Introduction

Liquid Chromatography coupled to Mass Spectrometry (LC-MS) has emerged as a leading tool in metabolomics as it provides both high sensitivity and high throughput. Much work has been done to produce robust experimental methods but data analysis bottlenecks still exist. The identification of known and unknown compounds detected was recently highlighted as one such bottleneck (Johnson et al. 2016).

Metabolite identification is typically performed by comparing experimental MS/MS spectra and retention times against reference spectra acquired from authentic standards on the same LC-MS system, see the recommendations of the metabolomics standards initiative (Sumner et al. 2007). A set of reference MS/MS spectra, with associated metadata, is known as a spectral library. The large variety of mass spectrometry platforms, protocols, and biological systems still motivates metabolomics labs and core facilities to create their own spectral libraries, especially when metabolite identification of a high confidence is required.

Generating spectral libraries is costly both due to the high price of standards and the effort of manual data curation needed for the highest quality libraries (Kind et al. 2009). Several initiatives exist to gather reference spectra as community resources, including <u>GNPS</u> (Wang et al. 2016), <u>European</u> <u>MassBank</u> (Stravs et al. 2013) and <u>MetaboLights</u> (Haug et al. 2013). Such large-scale efforts have the capability to integrate spectral libraries from different sources by crowdsourcing the contribution of individual annotated spectra. To enable the whole community to contribute, there is a strong need for software and standardized methods for creating, curating, and sharing a spectral library.

Here, we developed an open-source web application *curatr* for creating, curating, and sharing a custom spectral library. Curatr is complementary to mass spectrometry visualization tools, spectral matching software, and online data analysis and sharing platforms and repositories. Curatr stores information about standards analyzed, streamlines extraction of candidate MS/MS spectra, offers a graphical user interface to facilitate manual curation of MS/MS spectra, offers a searchable catalogue of all spectra contained in the library, and provides options for exporting the library to be used in commercial or open-source software as well as for submission to external spectral repositories.

### Description

Curatr is a web application that consists of a database, a server and a web interface. The application is written in Python using the Django framework. The open source project code and documentation is available at http://github.com/alexandrovteam/curatr, where issues can be reported. The server deployment has been tested under Linux (Ubuntu 14.04.4). User interaction is performed through the web interface that can be accessed from any computer using a modern web browser. All visitors can view and export the library but only signed-in users have the ability to upload and curate datasets.

Curatr has been designed to accept data in the mzML format from any full-scan MS system including time of flight (TOF), quadrupole TOF (qTOF), or Fourier transform (Orbitrap, FT), analyzers coupled to either liquid-chromatography or direct infusion sample inlets. The software was tested on LC-MS/MS datasets collected on an Infinity 1260 (Agilent, Waldbronn, Germany) LC and a Vanquish UHPLC systems (ThermoFisher Scientific, Bremen, Germany) coupled to a Q-Exactive Plus mass spectrometer (ThermoFisher Scientific). Before uploading to curatr, centroided data was exported to the mzML format using MSConvert (ProteoWizard, (Kessner et al. 2008)).

### Curating a spectral library

The steps of curation are shown schematically in Figure 1: Authentic standards are obtained and molecular information along with their provenance documented in curatr. For a standard, curatr only requires the name and molecular formula of the molecule it represents but we encourage also supplying its InChi code and PubChem ID. Standard provenance,such as vendor and catalogue number can also be recorded. A (LC)-MS/MS data set is then acquired, to increase throughput we recommend multiplexing standards. Following acquisition a user submits the data set through the web interface detailing: which standards were analysed; possible adducts; and details on the experimental conditions. Curatr then automatically extracts ion chromatograms and candidate fragmentation spectra for each standard-adduct ion based on a user-specified precursor mass tolerance (in ppm). As processing is performed asynchronously on the server, multiple datasets can be submitted.The raw data uploaded to curatr is stored in one place that simplifies the archiving.

**Figure 1.**
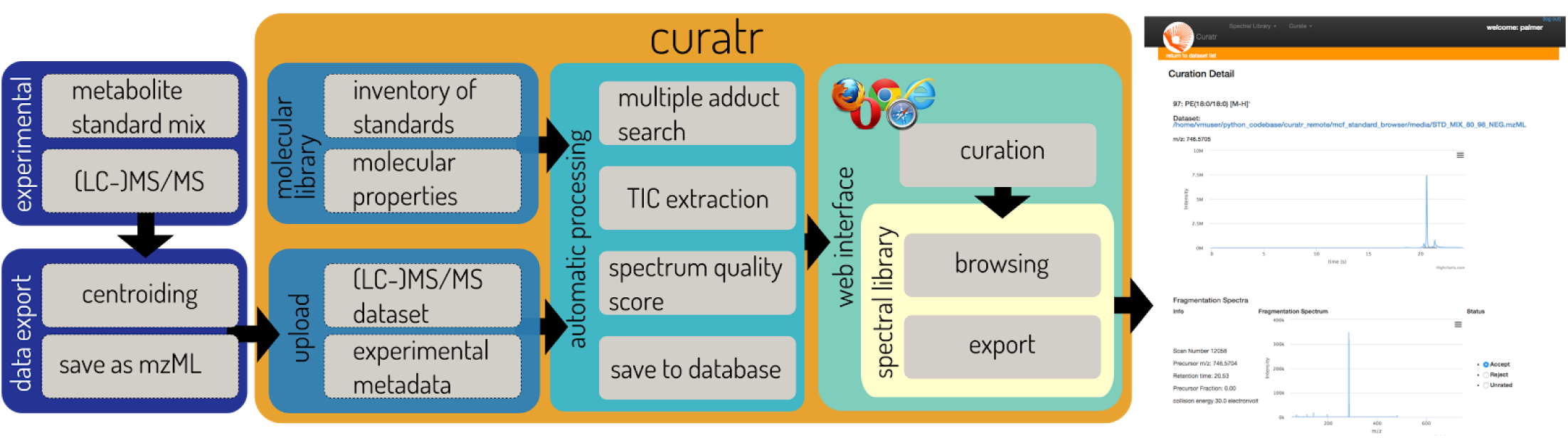
The workflow for creating and curating a spectral library using curatr. A user acquires authentic standards; enters their provenance and molecular information into curatr; performs LC-MS on a mixture of standards; uploads the dataset to curatr; and selects the best spectra offered within curatr to be assigned to a standard. Experimental sections are shown in purple and curation with curatr is contained in the orange region. The web-browser screenshot shows an example of selecting a spectrum for the library, a user is shown the XIC and associated MS/MS for the compound.

Once processed, a dataset can be selected from within the web interface. The web interface enables the curation to be performed from any computer and operating system. Users are shown candidate fragmentation spectra for each standard from which representative spectra can be selected for the spectral library. To aid in the selection, users can simultaneously view the extracted ion chromatogram and the tandem spectra. Additionally, we report a ‘precursor fraction’ metric that we have found to be useful to quantify the amount of possible interference from multiple peaks being present in the isolation window and consequently fragmented together. We defined the precursor fraction as the intensity of the selected precursor ion divided by the total intensity of the peaks within the quadrupole isolation window. A value close to 1.0 indicates that there is purely the intended precursor present. Whilst it is likely to be instrument dependent we have found that a value less than 0.5 (50%) indicates that additional peaks may be present and the fragmentation spectrum should be carefully inspected for interference. A user can select one or more spectra to be representative of the standard and add them to the spectral library.

### Exploring and exporting the spectral library

The primary browsing mechanism within the web interface is the molecular library, where any standards of the same molecule are grouped together. This library can be searched by molecular name, formula or m/z of any adduct. The curated fragmentation spectra of any molecule can be viewed on a single page allowing comparison across adducts, fragmentation energies, or between standards of the same molecule. Each spectrum is assigned a SPLASH spectral identifier (Wohlgemuth et al. 2016).

All molecules and every individual spectrum in the library can be directly hyperlinked for simple sharing. Additionally, the spectral library can be exported into plain text format (.tsv); mass spectrometry formats MGF and MSP; and additional custom formats. The selection currently implemented allows users to load the library into spectral matching software e.g. Progenesis QI (Waters, Wilmslow, UK) as well as to submit to online collections of reference spectra and spectral libraries e.g. GNPS, European MassBank and MetaboLights.

Curatr has been used to create and curate the EMBL Metabolomics Core Facility spectral library, an open-access library of LC-Orbitrap-MS/MS spectra from over 450 standards, and to share it publicly at http://curatr.mcf.embl.de/.

## Conclusion

Curatr is an open source web application for creation, curation, and sharing of a mass spectrometry spectral library. Our key ambition was to support individual labs in doing these tasks and furthermore expand the capabilities of the metabolomics community through streamlining sharing and exchange of individual libraries.

## Acknowledgements

We acknowledge resources from the EMBL Metabolomics Core Facility that were instrumental to test and improve the curatr software, support of the EMBL IT Services for deployment of the curatr, as well as funding from the European Union’s Horizon2020 programme under the grant agreement No. 634402.

